# Occurrence of the Humboldt Penguin (*Spheniscus humboldti*, Meyen, 1834) in El Ferrol Bay, Chimbote, Peru, and strategies for its conservation

**DOI:** 10.1101/2024.12.10.627413

**Authors:** Rómulo E. Loayza-Aguilar, Heillyn Medina-Quezada, Fernando Merino, Guillermo B. Saldaña-Rojas, Sorayda Mendoza, Gustavo E. Olivos-Ramirez

## Abstract

The Humboldt penguin is an emblematic species of the Humboldt ecosystem of Peru and Chile, declared as vulnerable in The IUCN Red List of Threatened Species in 2020. By 2023, its population was estimated around 9,900 pairs, which means a decrease of 93% of its population in 70 years. In response to a warning of its occurrence on the Guaneras Islands located in El Ferrol Bay (Chimbote, Peru), the objective of this study was to verify its presence *in situ*. Between October and November 2024, four expeditions were carried out on the leeward side of the Islote Peña Blanca (White Rock Islet), Isla Blanca (White Island), Isla Ferrol del Norte (Northern Ferrol Island), Isla Ferrol del Centro (Central Ferrol Island), and Isla Ferrol del Sur (Southern Ferrol Island). The presence of 149 individuals, 5 of them juveniles, and one chick accompanied by its parent, was verified on Northern Ferrol Island, Central Ferrol Island, and Southern Ferrol Island. It is estimated that the presence of these organisms is explained by the improvement of oceanographic conditions in this bay. Considering the high vulnerability of the species, the local community must articulate efforts to guarantee the establishment of its population in this particular habitat.

## INTRODUCTION

Peru ranks as the second most biodiverse country in the world for bird species, accounting for 1,860 species (Ugarte *et al*., 2023). However, this extraordinary diversity is not accompanied by exhaustive scientific knowledge. In many cases, critical aspects such as species distribution, population size, ecology, reproductive rates, chick survival, juvenile mortality, diet, seasonal movements, and habitat requirements remain poorly understood or entirely unknown (Angulo, 2018; McGill *et al*., 2021).

*Spheniscus humboldti,* commonly known as the “Humboldt penguin” or “baby bird” (Murphy, 1936), is a species that deserves significant attention from bird researchers and conservation managers due to its status as an emblematic component of the Humboldt ecosystem. This species is endemic to the Pacific Ocean along the western coast of South America, with a range extending from Isla Foca (Seal Island) in Peru (5°12′S) to Corral in Chile (39°52′S) (SERFOR, 2018; McGill *et al*., 2021). The Humboldt penguin is classified as vulnerable on the IUCN Red List of Threatened Species, with a declining population (McGill et al., 2021). Furthermore, the Peruvian government has officially designated it as an Endangered Species (Decreto Supremo 004-2014-MINAGRI; SERFOR, 2018).

This classification reflects significant challenges in preserving the species, as its population dynamics are of great concern. Historical counts from the late 1850s estimated approximately 1 million individuals, a number that plummeted to 9,000 in 2014 (Angulo, 2018). In 2019, studies recorded fewer than 10,000 individuals (McGill *et al*., 2021). These alarming statistics underscore the urgent need for conservation efforts.

According to Simeone (1996), Simeone *et al*. (1998), Simeone *et al*. (2002, 2003), Simeone & Luna-Jorquera (2012), Luna (2016), Cortés-Hinojosa *et al*. (2017), Luna-Jorquera *et al*. (2017), Angulo (2018), Sepúlveda *et al*. (2018), SERFOR (2018), Simeone *et al*. (2018), and McGill *et al*. (2021), the underlying causes of risk explaining the threatened extinction of *S*. *humboldti* are diverse. 1) The decline of areas for their reproduction on islands and at the same time the substrate (hardened guano where they build their breeding nests) due to the unplanned, irrational and illegal extraction of guano. 2) By incidental fishing with curtain nets and seines. 3) By intentional action through the removal of eggs, illegal capture of adults for use as pets, exhibition in zoos, use of their skin, or human food. 4) Disturbance of their sleeping, resting, and breeding areas by unplanned tourism, since this animal is easily perturbed by human presence, causing stress, especially during the breeding period. 5) Contamination by plastics, hydrocarbon spills and oils that threaten their health, which could be a direct cause of death. 6) Competition with *Homo sapiens* for *Engraulis ringens* (anchovy), which is their main food item. 7) Indirectly, the presence of El Niño events, since this phenomenon is directly related to the decrease in anchovy population. Facing this problem, in the short term, it is urgent to increase the level of collaboration among the scientific community to find and propose alternatives that allow the sustainable management of the populations, and in the medium and long term, to educate and raise public awareness on how they can help in the conservation of the Humboldt penguin (McGill et al., 2021).

The presence of *S*. *humboldti* in the Ferrol Islands within El Ferrol Bay was brought to light through accounts from a group of artisanal fishermen. This discovery raised significant interest, as the species had not been observed in the area since 1988, and no scientific data had been available to confirm its presence. Historical Humboldt penguin censuses during the moulting season from 1999 to 2010 documented the species in Huarmey (120 km south of El Ferrol Bay) in 2010. Subsequently, during censuses conducted from 2011 to 2020, 42 individuals were observed in 2018 on Santa Island, located 7 km northwest of the White Island (McGill et al., 2021). These findings prompted the authors to investigate and verify the reports *in situ*.

In this sense, the purpose of this work was to verify and document the presence of the Humboldt penguin in El Ferrol Bay, Chimbote, Peru. This research aims to provide valuable scientific data to support the efforts of competent authorities in the sustainable management of Peruvian wildlife. Additionally, it seeks to complement the initiatives of local government agencies, universities, researchers, and other institutions engaged in conserving diversity. Moreover, by addressing anthropogenic pressures, this study aspires to contribute to the stabilization and growth of the Humboldt penguin population within the islands of El Ferrol Bay in the short term.

## METHODS

### *In situ* observation approach

Between October and November 2024, four surveys were conducted between 07:00 and 13:00 hours aboard a small motorized boat. The surveys focused on the leeward side of Peña Blanca Islet, White Island, Northern Ferrol Island, Central Ferrol Island, and Southern Ferrol Island. Using binoculars (10 x 42) and a photographic camera, the presence of *S*. *humboldti* was recorded and photographed. To count the individuals, the photographs were analyzed on a large-screen computer. Individuals were categorized by life stage as chicks, juveniles, or adults. During the surveys, environmental parameters such as cloud cover, wind speed (measured with an anemometer), air temperature, and water temperature (measured with a simple thermometer) were also recorded.

## RESULTS

### Ecological and Geographic Features

El Ferrol Bay is located in the province of Santa, Ancash, Peru (Figure 1A), between coordinates 09°04’11” South Latitude and 078°35’31.8” West Longitude. Its coastline encompasses the districts of Chimbote and Nuevo Chimbote (Figure 1B) (Loayza, 2021). The environment is characterized by a temperate desert climate, with an average air temperature of 21.0°C (maximum 24.1°C, minimum 16.1°C) and a rainfall level ranging from 0.9 to 1.85 mm per month.

**Figure 1.**
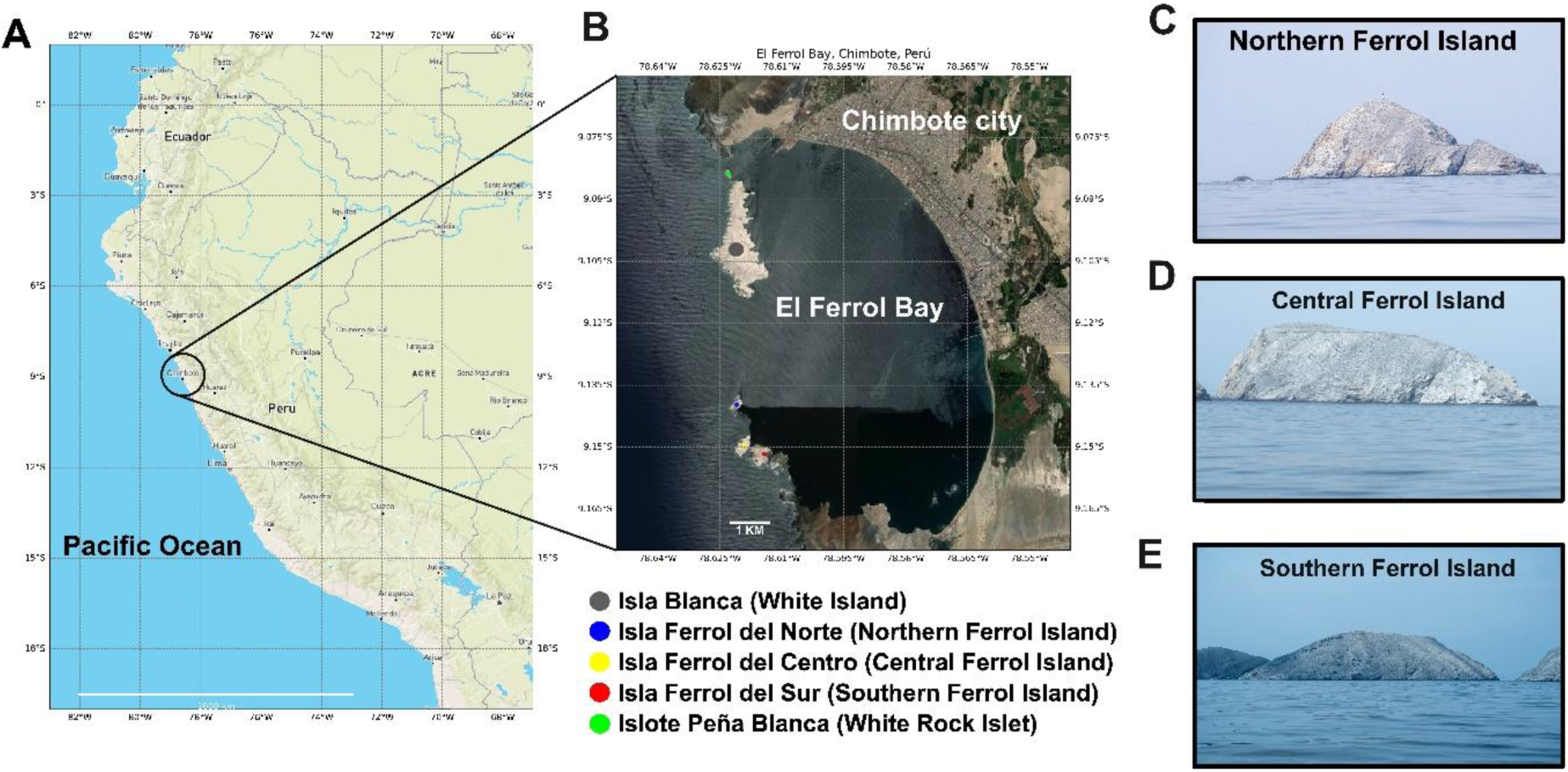
Geographical location of El Ferrol Bay, Chimbote, Peru. A) Geographical map of Peru. B) Geographical location of El Ferrol Bay and its islands. Panoramic view of the North (C), Center (D) and South (E) Ferrol Islands.

The bay is semi-enclosed, with a length of 11.19 km, a width of 6.57 km, and a coastline of 15 km (DHN, 2003), covering an area of 75.518 km² (Tresierra *et al*., 2007). It is defined by the Península Hill to the south and La Paz Hill to the north, along with the presence of four islands: Southern Ferrol, Central Ferrol, Northern Ferrol (Figure 1C-E), ranging from 84 to 123 meters in altitude, and the White Island, which reaches 204 meters. Between La Paz Hill and the White Island lies the White Rock Islet, with an altitude of 20 meters and a length of approximately 240 meters. Together, these orographic features protect the bay from winds, swells, and tsunamis, creating three channels that connect the bay to the open sea from south to north: the Ferrol Passage (760 m), the Middle Passage (2,700 m), and the Northern Passage (560 m) (Loayza, 2020). These islands feature several caves, most notably on the White Island. The tides are semi-diurnal, with an amplitude of 0.7 m, reaching 0.94 m during spring tides (Teves, 1999), generating currents with maximum speeds of 15 cm s⁻¹ in the Middle Passage. However, the average current speeds are low compared to other bays, which typically have average speeds greater than 30 cm s⁻¹ (Guzmán, 2006). Winds, ranging from 5 to 8.1 m s⁻¹ (Guzmán, 2014), with occasional gusts reaching 15-17 m s⁻¹ (Teves, 1999), have minimal influence on the generation of currents in the bay, leading to slow circulation with average current speeds between 6.5 and 14 cm s⁻¹, and a maximum speed of 38 cm s⁻¹ (Guzmán, 2014).

The four previously mentioned islands are desert-like and are characterized by the absence of fresh water and terrestrial vegetation. These islands are rich in guano (Figure 2A), with the White Island in particular serving as a significant habitat for seabirds. The most notable species in terms of population numbers include *Sula nebouxii* (Camanay), *Sula variegata* (Blue-footed Booby), *Phalacrocorax bougainvillii* (Guanay Cormorant), *Pelecanus thagus* (Peruvian Pelican), *Phalacrocorax gaimardi* (Red-legged Cormorant), and *Larosterna inca* (Inca Tern). Two of these islands feature remnants of infrastructure used for guano extraction (Figure 2D), dating back to the period known as the “Guano Fever,” which began in the mid-19th century (Méndez, 1987). On these islands, the *Microlophus peruvianus* lizard was introduced during the guano extraction period to control the tick population that posed a threat to the workers.

**Figure 2.**
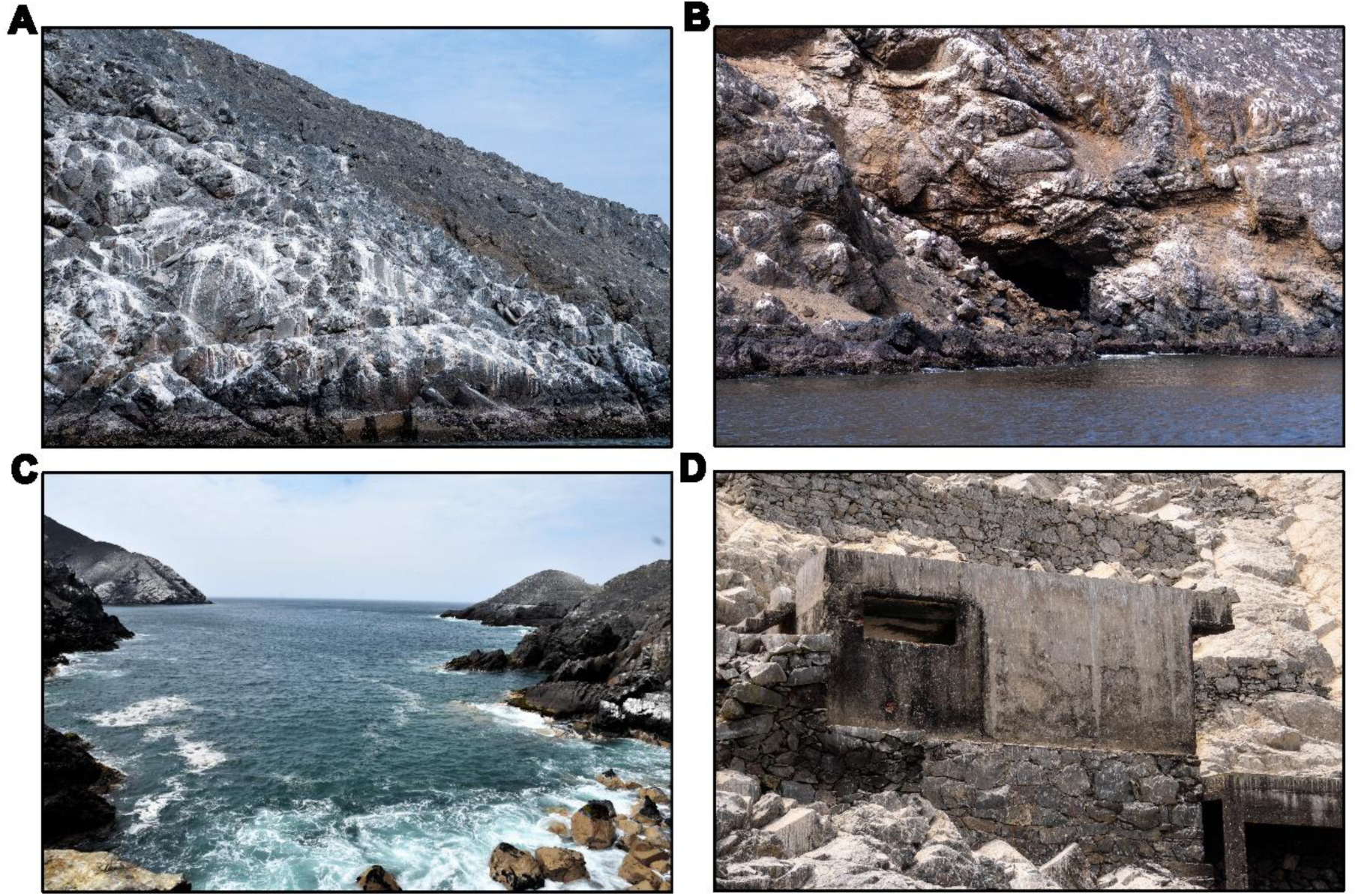
A) Guano deposits on the South Ferrol Island. B) Cave located on the South Ferrol Island. C) View of the water quality at the South Ferrol Island. D) Guano extraction infrastructure dating from the late 1800s.

### Occurrence of the Humboldt Penguin

During the surveys, *S*. *humboldti* was recorded on all three Ferrol Islands, but not on Peña Blanca Islet or White Island. All individuals were found near the intertidal zone (Figure 3D). A total of 149 individuals were counted, including 5 juveniles (Figure 3B) and one chick accompanied by its parent (Figure 3A). Some of them inhabit in couples (Figure 1C). During the surveys, sky coverage ranged from 8/8 to 1/8, air velocity averaged 2.4 ± 1.61 m s⁻¹, air temperature was 19.44 ± 2.23 °C, and water temperature was 16.4 ± 0.96 °C.

**Figure 3.**
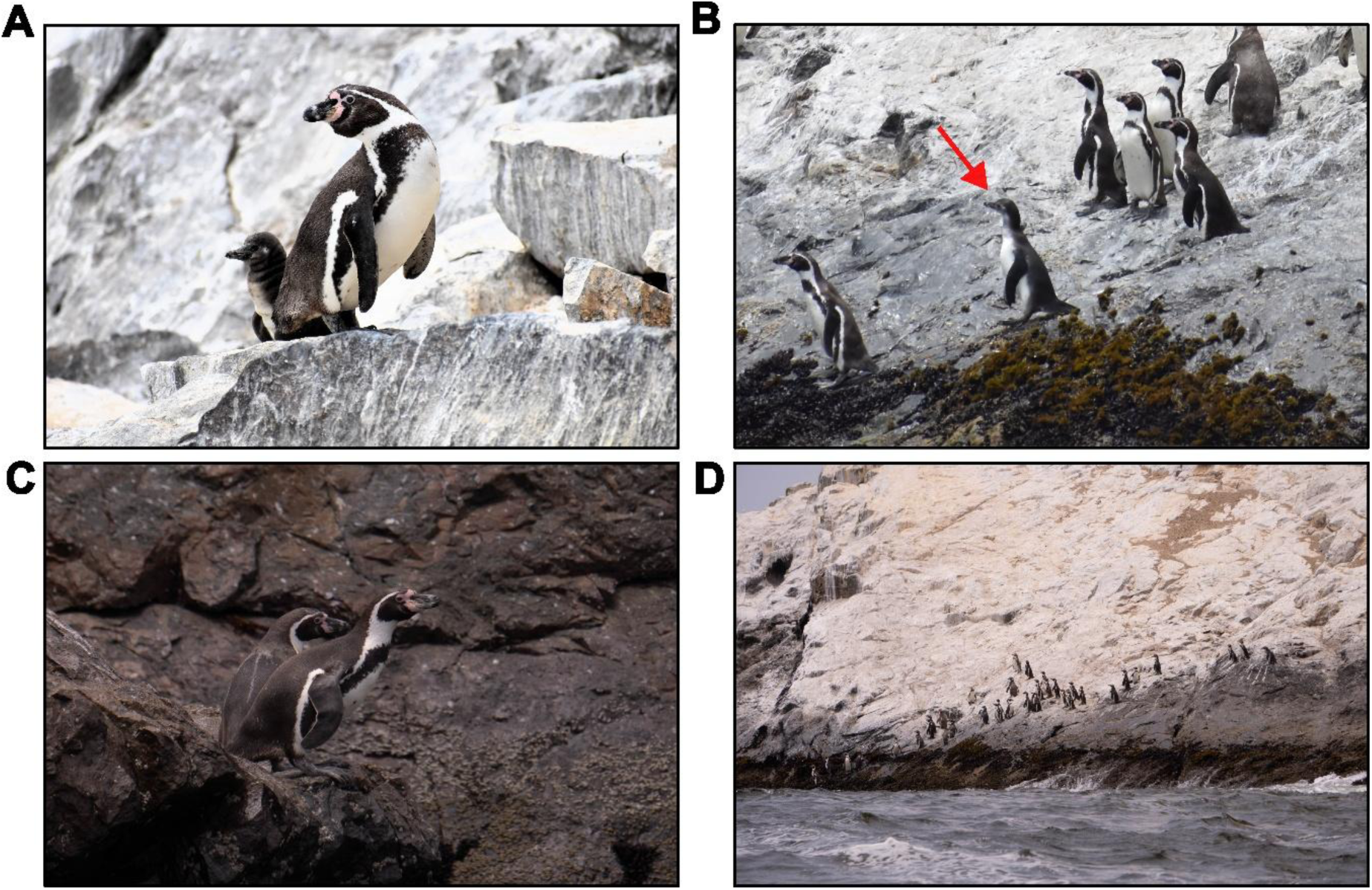
Occurrence of the Humboldt Penguin in the Ferrol islands. A) Chick accompanied by its parent. B) Shows the presence of a juvenile (red arrow). C) Pair of adults. D) Shows a colony of 37 penguins settled on South Ferrol Island.

## DISCUSSION

### Factors explaining the population settlement

The presence of *S. humboldti* in El Ferrol Bay could indicate improvements in its physical, chemical, biological, and sedimentological conditions, as the ecosystem is in the process of recovery (Figure 2C) after having been a receptor of effluents from the fishing industry between 1954 and 2015, and from the steel industry between 1958 and 2012. In addition to this industrial impact, domestic effluents from a rapidly growing population, now exceeding 250,000 peopcavele, were also introduced (Loayza, 2022). What is critical about this pollution process is that all the aforementioned effluents were discharged untreated into El Ferrol Bay, characterized by protein and lipid materials from 48 fishmeal and oil factories, inorganic material from the steel industry, and human fecal waste, as well as the disposal of all types of solid waste and hydrocarbons from the intense port activity. This process was further exacerbated by the slow circulation of the bay’s waters (cf. Guillen et al., 1998, UNS, 2000, Loayza, 1998, 2020a,b, 2021, MINAM, 2011, 2012).

Between the periods of 1994-1996 and 2002-2005, during fishmeal and oil production, dissolved oxygen concentrations were recorded between 0.0- and 0.73-ml l⁻¹ at the surface, oils and fats ranged from 0.65 to 20.78 mg l⁻¹, and biochemical oxygen demand (BOD₅) was greater than 10 mg O₂ l⁻¹, with peaks between 254 and 925 mg O₂ l⁻¹. Total coliform values ranged from 4.3 × 10² to 4.3 × 10³ NMP ml⁻¹, thermotolerant coliforms from 5 × 10³ to 4 × 10⁴ NMP ml⁻¹, and sulfide concentrations at the bottom exceeded 1 µg-at H₂S-S l⁻¹ (Orozco *et al*., 1997, Guillen *et al*., 1998, Tresierra *et al*., 2007, Sánchez *et al*., 2008). Additionally, an accumulation of 54,705,671 m³ of organic sludge was found at the bottom (Guzmán *et al*., 2002). In 2012, the steel industry ceased its pollution, and the fishing industry followed suit in 2015. However, the treatment of domestic effluents, which continue to be discharged untreated into the bay, and hydrocarbon pollution from vehicle emissions, primarily to the north of the bay, remain unresolved (Loayza, 2022).

Water contamination in El Ferrol Bay, caused by lipids and hydrocarbons on the surface or by the formation of emulsions, affected the plumage of birds and the gill systems of fish. The high biochemical oxygen demand generated hypoxic and generally anoxic conditions, directly impacting aerobic organisms. Water transparency decreased, negatively affecting the photosynthetic rate of phytoplankton. The production of toxic sulfurous gases, resulting from anaerobic degradation at the bottom, impacted the water column. These conditions likely constituted “barriers” preventing the Humboldt penguin (*S*. *humboldti*) from inhabiting El Ferrol Bay. This assertion is supported by McGill et al. (2021), who report the presence of this species between 2009-2018 in Huarmey (120 km south of the bay), in 2018 at Santa Island (7 km northwest of Isla Blanca), and in 2015 at Isla Chao and 2018 at Isla Corcovado, both located further north (Fig. 3). All of these ecosystems were free from the issues described for El Ferrol Bay.

In this context, it is highly likely that the species was present in El Ferrol Bay before the period of disturbance mentioned, as Peña Blanca Islet and the four guano-rich islands (Banco Centra de Reserva, 1962) would have provided an ideal habitat for population development. These islands offered abundant hardened guano (Figure 2A), caves (Figure 2B), cliffs, protection from waves, and an abundance of food, such as *E*. *ringens* and *Odontestes regia*, which are its primary food sources (Wilson *et al*., 1995).

Additionally, it can be inferred that the population of *S. humboldti* in El Ferrol Bay likely experienced two negative phases. First, the exploitation of guano from the islands in El Ferrol Bay, which took place from the mid-19th century (Duffy, 1994) until the early 1900s, decimated the guano layers, leaving a very thin layer that was insufficient for nesting and reproduction. This likely forced the penguins to abandon the habitat (Murphy, 1936). The second impact was caused by industrial and domestic effluents, as well as hydrocarbons and solid waste from port activities, beginning in 1954. However, these factors have decreased significantly in the past decade. Over time, these negative factors likely acted in a cascading manner, preventing the population of this species from maintaining itself in the bay.

### Conservation strategies

The current situation presented by the presence of *S*. *humboldti* on the three Ferrol Islands in El Ferrol Bay represents a serious commitment for the competent authorities, academia, benefactor institutions, and civil society. It is essential to coordinate efforts and establish strategies that will allow the species to consolidate its population on the islands of El Ferrol Bay.

For the development of strategies aimed at encouraging the consolidation of *S. humboldti* in El Ferrol Bay, it is important to consider that this species has evolved its wings into flippers, thus losing the ability to fly (Schweigger, 1964), making it vulnerable to poaching. Consequently, they require underground galleries in the hardened and accumulated guano, where they live, lay, and incubate their eggs (Murphy, 1936). Hardened guano allows the construction of burrow nests, which protect the chicks from dehydration and predation by seagulls (Hays, 1983; Duffy, 1990). In this context, the lack of substrate for nest construction and the reduction of optimal nesting sites lead to a decrease in reproductive success (Paredes & Zavalaga, 2001; Paredes et al., 2003).

Furthermore, most of these organisms exhibit activity within a 30 km radius of their colonies and at depths of less than 30 meters (Boersma *et al*., 2007; Mella, 2020). As a result, they are highly susceptible to becoming entangled in fishing nets (Taylor *et al*., 2004). Their plumage undergoes a complete molt, during which they do not enter the water or feed for a period of 10 to 12 days, making them particularly vulnerable to predators such as sea lions, otters, seagulls, and poaching. They are also exposed to stress from unplanned visits related to recreational and tourism activities. Reproduction begins at around three years of age (September to April), with the females laying two eggs. The incubation period lasts for 40 days, and the chicks depend on their parents for care until they reach three months of age (Zavalaga & Paredes, 1997; Paredes *et al*., 2002).

The presence of *S*. *humboldti* in El Ferrol Bay is significant, as it is an emblematic species of the Peruvian coast, regarded as the most charismatic bird among those inhabiting the Peruvian shoreline (Schweigger, 1964). However, it has been declared Endangered (Decreto Supremo 004-2014-MINAGRI), which demands urgent action, similar to efforts made in other regions (ACOREMA, 2013; Boersma *et al*., 2019). To address this, several measures should be implemented:

a. A research and permanent monitoring program for the species population. This should follow recommendations from UNEP-WCMC (2003), McGill *et al*. (2021), and Amaro *et al*. (2024), including addressing the impacts of both regulated and unregulated tourism.
b. A sustained environmental education program (UNEP-WCMC, 2003) to raise awareness and foster conservation. This program should focus on educating formal basic and higher education institutions, authorities (local government, Directorate of Regional Production, Environmental Evaluation and Inspection Agency, Harbor Master), the general public, and stakeholders involved in the bay (artisanal fishermen, shipowners, canned food and fishmeal factories, water management companies, PETROPERÚ, and tourism operators).
c. Implement measures to support the species’ population establishment on the islands. One example is the construction of artificial nests, as suggested by Simeone *et al*. (2010). This model, used successfully with African penguins on the Namibian coast, involves using 120-liter plastic containers, as shown in Figure 4A and B. This measure would reduce predation, especially by birds, and improve reproductive success.

**Figure 4.**
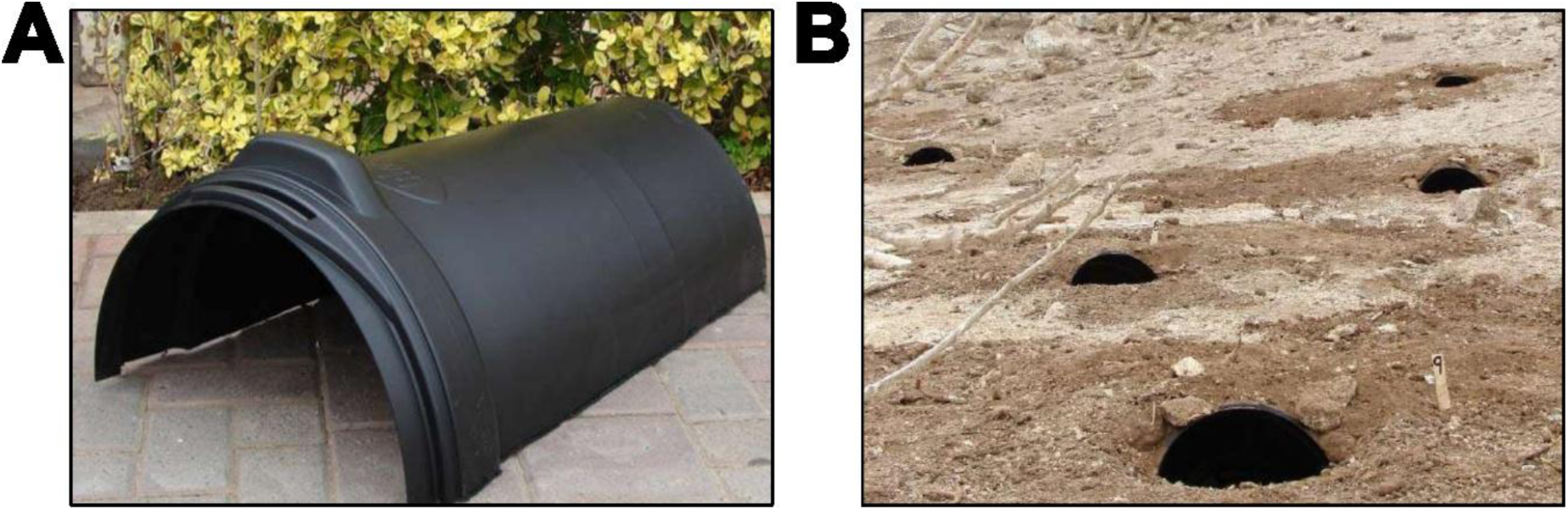
A) y B) Strategy for the construction of artificial nests for *S*. *humboldti* (Image taken from Simeone et al., 2010).

From a political and legal perspective:

a. A plan should be established to disseminate the legal and regulatory framework that protects the species, with the aim of reducing both targeted and incidental capture (Zavalaga & Alfaro, 2015).
b. Agreements should be reached between the competent authorities and fishing unions to establish fishing restriction zones and define the types of nets to be used (UNEP-WCMC, 2003) around the islands in El Ferrol Bay.
c. Agreements should be established with the owners of vessels (small, medium, and large) to prevent the introduction of invasive species such as rats, cats, and dogs, as they are predators that could significantly affect the *S*. *humboldti* populations (Simeone *et al*., 2003; Simeone & Luna-Jorquera, 2012; Sepúlveda *et al*., 2018).
d. Vessel operators in the northern parking zone of the bay (approximately 700 boats) should be encouraged to avoid the disposal of solid waste, particularly plastics, as these could affect *S*. *humboldti* through accidental or intentional ingestion, leading to choking, or exposure to microorganisms and heavy metals attached to the plastic surface. This damage could also extend to chicks, as they may ingest food contaminated with plastic debris (ACOREMA, 2013).
e. The potential extraction of guano from these islands should be avoided, allowing the guano layer to grow sufficiently for these organisms to build their burrows, which are essential for the consolidation of the population in this ecosystem.
f. Efforts should be made to ensure that the four islands and the islet in El Ferrol Bay are included in the National Reserve of the System of Islands, Islets, and Guano Points, established in 2009 by Supreme Decree 024-2009-MINAM.

### Conclusions

The presence of *S. humboldti* on the guano islands of El Ferrol, located in El Ferrol Bay, may indicate that the bay currently offers suitable conditions for the establishment of this species. This follows a period of over 50 years during which the bay suffered severe impacts due to the uncontrolled growth of the fishing industry (fishmeal and oil), which discharged its untreated effluents into the bay. Since the Humboldt Penguin is an emblematic species of the Humboldt ecosystem, yet considered “Vulnerable” and in decline by the IUCN Red List, as well as “Endangered” by the Peruvian government, its presence on the three Ferrol Islands in El Ferrol Bay grants an excellent opportunity to implement protective measures for the species. These measures should aim to consolidate its population. To achieve this, efforts must be coordinated among the relevant authorities, universities, NGOs, tour operators, fishing guilds, professional associations, and the general public.

## Acknowledgments

To Mr. Freddy Arteaga, artisanal fisherman from the port of Chimbote, for sharing information about the presence of the Humboldt Penguin in El Ferrol Bay. To the School of Biology in Aquaculture at the National University of Santa for facilitating visits to El Ferrol Bay. To students Yohan Arbildo Rodríguez and Steven Espinola Alva, both from the School of Biology in Aquaculture at the Universidad Nacional del Santa, for sharing some photographs. To the anonymous reviewers of this document for their valuable contributions.

## Authors’ contribution

Rómulo Loayza: Formal analysis, Conceptualization, Investigation, Methodology, Writing-reviewing, Editing, Visualization. Heillyn Medina: Investigation, Methodology, Reviewing. Juan Merino: Methodology, Resources, Reviewing. Guillermo Saldaña: Methodology, Resources, Reviewing. Sorayda Mendoza: Methodology, Resources, Reviewing. Gustavo Olivos: Writing and Reviewing.

